# Host genetic regulation of fermentation-related cecal microbial taxa is associated with loin muscle deposition in meat rabbits

**DOI:** 10.64898/2026.06.09.731016

**Authors:** Yan Di, Qianli Fu, Kerui Xie, Zichen Song, Wenqiang Li, Xianyao Li, Qin Zhang, Chao Ning, Dan Wang, Xinzhong Fan

**Affiliations:** College of Animal Science and Technology, Shandong Agricultural University, Taian, 271017, China; College of Agriculture and Biology, Liaocheng University, Liaocheng 252059, China

**Author notes:** Corresponding author. Email address (Xinzhong Fan) Email address (Dan Wang).

**Keywords:** Meat rabbit, Loin muscle deposition, Cecal microbiota, Hindgut fermentation, Host-microbiome interaction, Segment-specific microbiome

## Abstract

Loin muscle weight is an important indicator of carcass yield in meat rabbit production, but the host genetic and intestinal microbial factors associated with its variation remain poorly understood. Because the cecum is the primary site of hindgut fermentation in rabbits, we integrated whole-genome resequencing, cecal transcriptome profiling, and cecal and rectal 16S rRNA sequencing data from 321 Kangda meat rabbits, with rectal microbiome data used as a downstream comparative reference. Compared with the rectum, the cecum contained a richer microbial community, with more ASVs and genera and significantly higher microbial diversity, whereas predicted metabolic functions were largely conserved between the two segments. Loin muscle weight showed moderate SNP-based heritability (h² = 0.39), and the cecal microbiome explained a smaller but detectable proportion of phenotypic variation (m² = 0.13). Multi-strategy microbial screening identified 19 candidate cecal genera associated with loin muscle weight, with Methanosphaera showing the strongest negative association. Host GWAS prioritized candidate loci near MEX3C and TCF4 on chromosome 10, and integration of cecal cis-eQTL and GWAS summary statistics further prioritized GJB3 as a candidate gene associated with loin muscle weight. Consistent with the cecum-centered model, host genetic relatedness was weakly but significantly correlated with cecal microbial similarity, whereas no such global association was observed for the rectal microbiome. Microbial GWAS and SMR analyses further prioritized a cecal MRAP2–Methanobrevibacter association as the main host-regulated microbial signal, while rectal analyses identified distinct segment-specific signals, including SULF1–Roseburia. These findings suggest that host genetic variation may be linked to loin muscle deposition partly through cecal gene expression and fermentation-related cecal microbial taxa, with the rectal microbiome providing comparative evidence for hindgut segment specificity. This study provides candidate host and microbial targets for future functional validation and microbiome-informed nutritional strategies to improve carcass traits in meat rabbits.

## 1. Introduction

The meat rabbit has become an important source of meat in the global livestock industry, thanks to its short growth cycle, high fecundity, and meat characteristics of high protein and low fat (Carneiro et al., 2011; Matthee et al., 2004). Since the meat rabbit is a monogastric hindgut-fermenting herbivore, the cecum functions as the core site of hindgut fermentation, while the rectum serves as a downstream intestinal segment for comparative reference. In intensive meat rabbit farming, loin muscle (LM) weight directly determines the yield of high-value meat cuts and is a core indicator for measuring economic returns. Therefore, elucidating the interactions among host genetics, cecal microbial communities, and their contributions to LM deposition is essential for improving growth performance and supporting both nutritional interventions and precision breeding in meat rabbits.

Previous studies have demonstrated a significant association between gut microbiota composition and host skeletal muscle development, a concept increasingly recognized as the “gut-muscle axis” (Lahiri et al., 2019). In hindgut-fermenting animals, the intestines are enriched with specific saccharolytic and butyrate-producing bacteria. These microorganisms efficiently ferment complex dietary polysaccharides to produce short-chain fatty acids (SCFAs) (Shah et al., 2019). Rather than merely providing supplementary energy to the host, these gut-derived metabolites act as critical signaling molecules that systemically drive skeletal muscle growth and protein deposition. Consistent with this physiological mechanism, recent studies utilizing germ-free animal models have confirmed that the absence of gut microbiota profoundly disrupts skeletal muscle morphogenesis and energy metabolism (Li et al., 2025). Similarly, specific microbial taxa have also been confirmed as critical drivers of skeletal muscle development across multiple species, including mice (Bárcena et al., 2019), cattle (Holman et al., 2024), and chickens (Peng et al., 2026).

Although dietary components (Cong et al., 2022) and environmental factors (Li et al., 2023) play important roles in shaping the gut microbiota, the host genetic background also exerts a decisive influence on the colonization and homeostasis maintenance of microbial communities. Acting as the host’s second genome, the gut microbiota co-evolves with the host and dynamically participates in energy harvesting and metabolic reactions (Bäckhed et al., 2004; Guarner and Malagelada, 2003). The gut microbiota was first proposed for genetic analysis as quantitative traits in 2010 (Benson et al., 2010). Subsequently, studies in humans (Goodrich et al., 2014), pigs (Chen et al., 2018a), and beef cattle (Li et al., 2019) have confirmed that the abundance of specific gut microbial taxa possesses moderate to high heritability. Through genome-wide association studies (GWAS), researchers have identified multiple host genetic loci that regulate microbial abundance. For instance, human LCT gene variations are significantly associated with *Bifidobacterium* abundance (Blekhman et al., 2015; Kurilshikov et al., 2021), while the ABO homologous locus in pigs affects the colonization of Erysipelotrichaceae (Yang et al., 2022). These findings suggest that the host may screen and enrich specific symbiotic microbial groups through particular genetic mechanisms to adapt to the requirements of hindgut fermentation.

However, current research on meat rabbits is largely confined to single phenotypic correlations or nutritional intervention analyses, and the biological mechanisms by which host genetic variation regulates cecal gene expression to shape microbial assembly and muscle deposition remain largely unclear. In meat rabbits, previous studies have observed differences in cecal microbiota among different body weight groups (Zeng et al., 2015). Although the latest study explored the effects of diet on growth and meat quality(Biasato et al., 2025), it also noted that a single nutritional strategy did not alter cecal microbiota stability; this suggests that more fundamental regulatory factors, particularly host genetic variation and its effects on cecal physiology, may underlie variation in microbial composition and production traits.

In this study, we systematically integrated host genome, cecal transcriptome, and cecal and rectal microbiome profiles from 321 meat rabbits to characterize the host genome–cecal transcriptome–microbiome axis associated with loin muscle weight. We aimed to identify host genetic variants, cecal gene-expression mediators and microbial taxa associated with loin muscle weight, and to elucidate potential biological pathways linking host genetics, cecal microbial communities, and muscle deposition in meat rabbits. These findings are expected to provide new insights into the biological regulation of complex production traits and offer a theoretical foundation for precision breeding and nutritional strategies targeting hindgut fermentation in meat rabbits.

In this study, we integrated host whole-genome resequencing, cecal transcriptome profiling, and cecal and rectal microbiome data from Kangda meat rabbits to investigate host genetic regulation of fermentation-related cecal microbial taxa associated with loin muscle deposition. Given the central role of the cecum in hindgut fermentation, we focused primarily on a cecum-centered host genome-cecal transcriptome-microbial taxon axis. Rectal microbiome data were included as a downstream comparative segment to evaluate whether host genetic associations with microbial taxa were shared across the hindgut or showed segment-specific patterns. These analyses aimed to identify candidate host variants, cecal gene-expression mediators, and microbial taxa associated with loin muscle deposition, thereby providing potential targets for future functional validation and microbiome-informed nutritional strategies in meat rabbits.

## 2. Materials and methods

### 2.1. Ethics statement

All experiments involving animals were conducted according to the ethical policies and procedures approved by the Institutional Animal Care and Use Committee of Shandong Agricultural University, China (SDAUA-2024-110).

### 2.2. Animals, phenotypic data and sample collection

This experiment used 321 healthy Kangda meat rabbits (116 females and 205 males) as the experimental sample, provided by the Kangda meat rabbit breeding farm in Huangdao District, Qingdao City, Shandong Province. The rabbits were randomly selected from two batches and ear-tagged. The breeding environment, immunization procedures and feed formulas remained consistent.

After the rabbits reached 70 days of age, they were fasted for 12 hours with free access to water prior to slaughter. A uniform slaughter method was applied, which involved stunning the rabbits electrically followed by bloodletting. The abdominal cavity was immediately opened using a scalpel, and the intestines were extracted. The cecum was then cut open using sterile surgical scissors to collect cecum tissue and cecal contents, followed by the collection of rectal contents from the distal rectum. Muscle samples were also collected simultaneously. All samples were rapidly frozen in liquid nitrogen and transferred to a −80°C freezer for storage. Loin muscle (LM) weight was recorded to the nearest 1 g using an electronic scale.

### 2.3. Study design and multi-omics framework

This study was designed to investigate host genetic regulation of fermentation-related cecal microbial taxa associated with loin muscle deposition in meat rabbits. Whole-genome resequencing data were used to characterize host genetic variation, cecal transcriptome data were used to identify host gene-expression mediators, and 16S rRNA sequencing data from cecal and rectal contents were used to characterize intestinal microbial communities. Because the cecum is the primary site of hindgut fermentation in rabbits and both cecal transcriptome and cecal microbiome data were available, the main analyses focused on a cecum-centered host genome-cecal transcriptome-microbial taxon axis. Rectal microbiome data were analyzed as a downstream comparative segment to evaluate whether host genetic associations with microbial taxa were shared across hindgut segments or showed segment-specific patterns. Accordingly, cecal microbiome analyses were used to identify microbial taxa associated with loin muscle weight, whereas rectal microbiome analyses were used mainly for comparative and segment-specific interpretation.

### 2.4. Whole-genome resequencing and Genotyping

A total of 321 muscle tissues were sent to Beijing Novogene Bioinformatics Technology Co., Ltd. and Beijing Biomarker Technologies Co., Ltd., where low-depth re-sequencing was performed using the BGI DNBSEQ-T7 and BGI MGISEQ-T7 sequencing platforms, respectively, to generate 150 bp paired-end reads. Adapter sequences and low-quality reads from the raw data were filtered using Fastp (version 0.19.6) (Chen et al., 2018c). Clean reads were mapped to the reference genome (GCF_964237555.1) using Bwa (v0.7.17) (Liu et al., 2016). Alignment results were sorted using Samtools (v1.9) (Li et al., 2009). Duplicate sequences were marked with GATK (v4.1.4.1) (McKenna et al., 2010), generating deduplicated BAM files, which were then indexed using Samtools. Genotyping was performed with BaseVar (v1.0) (Liu et al., 2024), and imputation was done using STITCH (Davies et al., 2016). Secondary imputation of the remaining missing sites was performed using Beagle (v5.4) (Browning and Browning, 2016). Finally, the obtained SNP data were filtered using PLINK (v1.90) (Purcell et al., 2007) with the following parameters: Hardy-Weinberg equilibrium (HWE) *P* > 10^-6^ and minor allele frequency□>□5%. A total of 20,961,519 SNPs were retained for subsequent analysis.

### 2.4. 16S rRNA gene sequencing and analysis

Microbial DNA from 321 cecal content samples and 112 rectal content samples was extracted using the standard CTAB method. The V3-V4 region of the intestinal microbiome was selected as the amplification target. PCR amplification was performed, followed by purification, quantification, and normalization of the amplicons to construct a sequencing library. The constructed library underwent quality control, and libraries that passed the quality check were sequenced using the Illumina HiSeq 2500. Filtered the raw reads using Trimmomatic (v0.33) (Bolger et al., 2014) to remove adapter sequences, barcode sequences, and low-quality sequences. Then, primer sequences were identified and removed using cutadapt (v1.9.1) (Marcel, 2011). Paired-end sequences were denoised and filtered using DADA2 (Callahan et al., 2016), retaining a length of 220 bp for both ends. We established a minimum overlap length of 4 base pairs. The high-quality clean reads were obtained for subsequent clustering analysis, which was performed by clustering the data into amplicon sequence variants (ASVs). A 100% similarity threshold was used for ASVs classification, with the Silva-138 database (Quast et al., 2013) for comparison. Taxonomic information was obtained for each classification level: Kingdom, Phylum, Class, Order, Family, Genus, and Species.

Use the phylogeny algorithm in Qiime2 (Bolyen et al., 2019) to cluster the effective data, the ASV feature table was rarefied to an even sequencing depth of 43,000 reads per sample. The Vegan package in R was used to calculate the Shannon index of α-diversity for microbiome samples from the cecum and rectum. Perform a unpaired Student’s t-test to assess the differences between the two. Picrust2 (Douglas et al., 2020) was employed to predict the potential functions of the microbiota in both intestines, comparing the KEGG database and MetaCyc database.For biological interpretation, fermentation-related microbial taxa were defined as taxa previously reported to participate in carbohydrate degradation, methanogenesis, short-chain fatty acid production, or hindgut fermentative metabolism, based on published literature and predicted microbial functional profiles.

### 2.5. Estimation of Host Heritability and Microbiability for Complex Traits

Genetic heritability of the phenotypes was calculated using GMAT (v1.2) (Wang et al., 2020) software. Sex and batch were included as covariates. The estimation model is as follows:

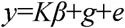

Here y represents the phenotype vector; β is the covariate vector (sex and batch); K is the corresponding coefficient matrix; e is the residual vector; and g is the effect vector of all SNPs. g is a vector of polygenic effects following the normal distribution N (0, Gσ^2^_g_), where G is the genetic relatedness matrix (GRM) calculated based on genome-wide marker information, σ^2^_g_ is the genetic variance (Yu et al., 2006). The GRM was constructed using the Gmatrix module in GMAT software, with the following formula:

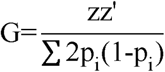

Where p_i_ represents the minor allele frequency at the i-th locus, and z is the marker matrix (coded as 0, 1, and 2), with each column subtracted from twice the minimum allele frequency.

After filtering for microbial genera with a detection rate > 30%, we used the relative abundances of microbial ASVs to construct the microbial relationship matrix (MRM) for calculating microbiability (*m^2^*). The microbial relationship matrix is constructed as follows:

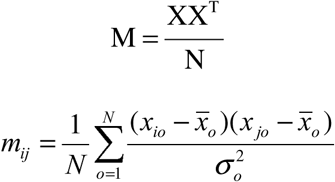

In the equation, M represents the MRM, X is the normalized ASVs matrix of size b×N, and m_ij_ is the refinement of M; x_io_ and x_jo_ represent the relative abundance of ASV o for individuals i and j. b is the number of rabbit used to estimate the MRM, and N is the number of ASVs used to estimate the MRM. The Z-score normalization formula for any element in Xs is as follows:

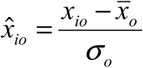

x_io_ represents the relative abundance of ASV o for individual i in gut segment s, and 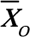 represents the average relative abundance of ASV o. σ_o_ represents the standard deviation of the relative abundance of ASV o.

Microbiability used the similar model as heritability, with g replaced by M to represent the effects of all ASVs. The GRM was replaced with MRM.

### 2.6. Genome-Wide Association Studies for Host Traits and Microbial Abundances

mGWAS (microbial Genome-Wide Association Study) used the univariate linear mixed model from the GMAT software, considering sex and batch effects as covariates. The model used is:

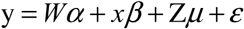

Here: y represents the vector of detection rate > 30% for microbial relative abundance at genus level. The detection rate between 30% and 60% was transformed into binary traits (0/1). For microbial genera with a detection rate exceeding 60%, zero observations were recoded as missing values. W represents the covariate matrix (including sex and batch effects). α represents the corresponding coefficients, including the intercept. x represents the vector of marker genotypes. β represents the effect size of the marker. μ represents the individual random effects following the normal distribution N (0, Gσ^2^_g_), where G is the genetic relatedness matrix. ε represents the residual vector.

Vector y in the aforementioned model parameters was then replaced with the vector of LM to perform GWAS. After filtering SNPs based on linkage disequilibrium (LD) using PLINK, we retained 457,229 SNPs for the cecum and 442,905 for the rectum. Therefore, the significance threshold for microbiota in the cecum was set to 1.09×10□□ (0.05/457,229), with the suggestive significance threshold being 2.19×10□^6^ (1/457,229). For the rectal contents microbiota, the significance threshold is set to 1.13×10□□ (0.05/442,905), and the suggestive significance threshold is 2.26×10□□ (1/442,905).

### 2.7. Transcriptome Analysis and cis-eQTL Mapping

Total RNA extracted from 112 cecum samples was evaluated for quality and integrity prior to library construction. Briefly, mRNA was isolated using Oligo (dT) magnetic beads and randomly fragmented. Double-stranded cDNA was synthesized using random hexamers, followed by purification, end repair, A-tailing, and adapter ligation. After size selection via AMPure XP beads, the library was generated by PCR amplification. Library quality, including concentration and insert size, was validated using Qubit 2.0, Agilent 2100, and Q-PCR. Sequencing was performed on the NovaSeq 6000 platform, generating 150 bp paired-end reads. Raw reads were quality-controlled using FastQC (v0.23.2) and aligned to the reference genome (GCF_964237555.1, mOryCun1.1) using STAR (v2.7.11b) (Dobin et al., 2013) in 2-pass mode. Gene-level read counts were quantified using featureCounts (v2.0.6) (Liao et al., 2014), and Transcripts Per Million (TPM) values were calculated. Genes with TPM < 0.1 or raw counts < 6 in more than 80% of the samples were excluded. Inter-sample expression variation was corrected using the Trimmed Mean of M-values (TMM) method in edgeR (Robinson et al., 2010), followed by a rank-based inverse normal transformation (INT) of the normalized expression.

*cis*-eQTL mapping was performed using a Linear Mixed Model (LMM) implemented in OmiGA (v1.0.3) (Teng et al., 2026). We tested SNPs located within a 1 Mb window upstream and downstream of the transcription start site (TSS) of each gene. Sex and 12 expression Principal Components (PCs) were selected as covariates. The number of PCs was automatically determined based on the rate of change in Percent Variance Explained (PVE). An eGene was defined as a gene with the corrected P-value across phenotypes < 0.05 and at least one variant with a P-value < variant-level significance threshold. This threshold was determined based on a 5% FDR level from 10,000 permutations.

### 2.8. Summary data-based Mendelian randomization analysis

The SMR test for pleiotropic association between the expression level of a gene and a complex trait of interest using summary-level data from GWAS and expression quantitative trait loci (eQTL) studies. We employed SMR software (v1.3.2) (Zhu et al., 2016) to integrate *cis*-eQTL summary data with summary statistics from host GWAS and mGWAS. The analysis was restricted to genomic regions with *P* < 10^-5^. For instrumental variable selection, under the prerequisite of identifying eGenes, significant *cis*-eQTLs were identified using the variant-level thresholds determined by the aforementioned permutation tests. To exclude false-positive associations resulting from linkage disequilibrium (LD) rather than causal relationship, we performed the HEIDI (heterogeneity in dependent instruments) test. A P_HEIDI > 0.05 indicated a lack of significant heterogeneity, supporting a potential causal relationship. Finally, the SMR P-values were corrected for multiple testing using the Benjamini-Hochberg (FDR) method to identify the final candidate association.

### 2.9. Statistical analysis

To identify microbial genera associated with host trait (LM), after removing genera with a detection rate < 30%, the Wilcoxon rank-sum test was used to assess the differences in trait between the top 20% and bottom 20% of rabbits based on the relative abundance of microbial genera. SRC (Spearman’s rank correlation) analysis was performed using the microbial relative abundances and traits. The Wilcoxon test and SRC P-values are corrected using FDR, with an adjusted *P* < 0.05 considered significant.

The detection rate between 30% and 60% were transformed into binary traits (0/1). For microbial genera with a detection rate exceeding 60%, zero observations were recoded as missing values, while the genera with a detection rate of less than 30% were removed. Perform multiple linear regression analysis for both types of variables in R, considering the effect of sex and batch. Then, use the StepAIC function in the MASS package to perform stepwise regression for both multiple linear regression results. The multiple linear regression model is:

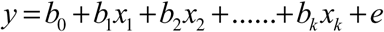

Here, y is the response variable (LM), b_0_ is the constant term, b_1_ to b_k_ are the coefficients of the independent variables, x is a vector of microbial genera, and e is the residual.

To identify gene expression associated with host trait (LM), Samples were categorized into high and low groups based on the top and bottom 20% of each trait. Before differential analysis, raw counts were filtered to retain only genes with a count greater than 10 in at least 20% of the samples. Differential expression analysis was then performed using the DESeq2 package. Differentially expressed genes (DEGs) were identified based on the criteria of *P* < 0.05 and |log_2_ FC| > 1. The TPM were first filtered to exclude low-abundance genes, retaining only those with a TPM > 0.5 in more than 50% of the samples. SRC were calculated between the filtered TPM and each host trait. The resulting P-values were adjusted for multiple testing using the Benjamini-Hochberg (FDR) method.

## 3. Results

### 3.1. Phenotypic and multi-omics data overview

The descriptive statistics for loin muscle (LM) weight were presented in Table S1. A Shapiro-Wilk test confirmed that LM followed a normal distribution in this experimental population (Additional file2 Fig. S1).

To obtain the host genomic variants, we generated an average of 12Gbp of clean bases from 321 rabbits, achieving an average sequencing depth of 2.10-fold. After stringent quality control, we identified a total of 20,961,519 SNPs. (Table S2).

We performed stringent quality control on the raw bulk RNA-sequencing data. A total of 784 G of raw data was generated, and after removing adapter sequences and low-quality reads, we obtained 760 G of clean data. The quality of the sequencing was high, as indicated by an average Q30 of 94.5%. The GC_Content was approximately 57%. The Unique_Mapping_Rate reached 86%, demonstrating the high efficiency of reads aligning to unique genomic locations (Table S3).

The raw data for the cecum consisted of 26,366,533 reads. After denoising and chimeric sequence removal, a total of 18,441,254 high-quality reads were obtained, with an average of 57,449 reads per sample. The raw data for the rectum consisted of 8,960,353 reads, with 6,823,590 clean reads, and an average of 60,924 reads per sample (Table S4). A total of 12,228 ASVs were identified in the cecum, classified into 30 phyla, 69 classes, 162 orders, 284 families, 572 genera, and 896 species. In the rectum, 6,586 ASVs were identified, classified into 25 phyla, 65 classes, 142 orders, 234 families, 424 genera, and 627 species. The proportion of clearly defined reads decreased at higher taxonomic levels, with 71.56% and 67.42% of reads confidently annotated at the species level in cecal and rectal samples, respectively. (Fig.S2). Overall, these data supported a cecum-centered multi-omics framework, with rectal microbiome data serving as a downstream comparative layer.

### 3.2. Cecal and rectal microbiomes show distinct community structures but conserved predicted metabolic functions

The dominant microbiota at phylum level in the cecum included: *Bacteroidota, Firmicutes*, *Verrucomicrobiota*, *Desulfobacterota*, and *Proteobacteria*. The relative abundance of these phyla was all above 1%. *Bacteroidota* and *Firmicutes* accounted for a total relative abundance of 88.8%. The relative abundance of the five dominant phyla accounted for 97.3%, and their detection rate was 100%. There were four dominant microbiota at phylum level overlap between the cecum and rectum. However, *Desulfobacterota* was replaced by *Cyanobacteria* in the rectum. *Firmicutes* and *Bacteroidota* accounted for a total relative abundance of 88.1% (Fig. 1A, Table S5). At the genus level, *Muribaculaceae* was the most abundant dominant genus in both cecum and rectum, reaching 18.3% and 24.3%, respectively (Fig. 1B, Table S6). The Shannon index was used to represent the α-diversity of the microbiota. The microbial diversity in the cecum was significantly higher than that in the rectum (*P* < 0.001) (Fig. 1C), indicating that although there was some overlap in the microbial communities between the two segments, the microbial species in the cecum were significantly higher than those in the rectum.

**Fig. 1.**
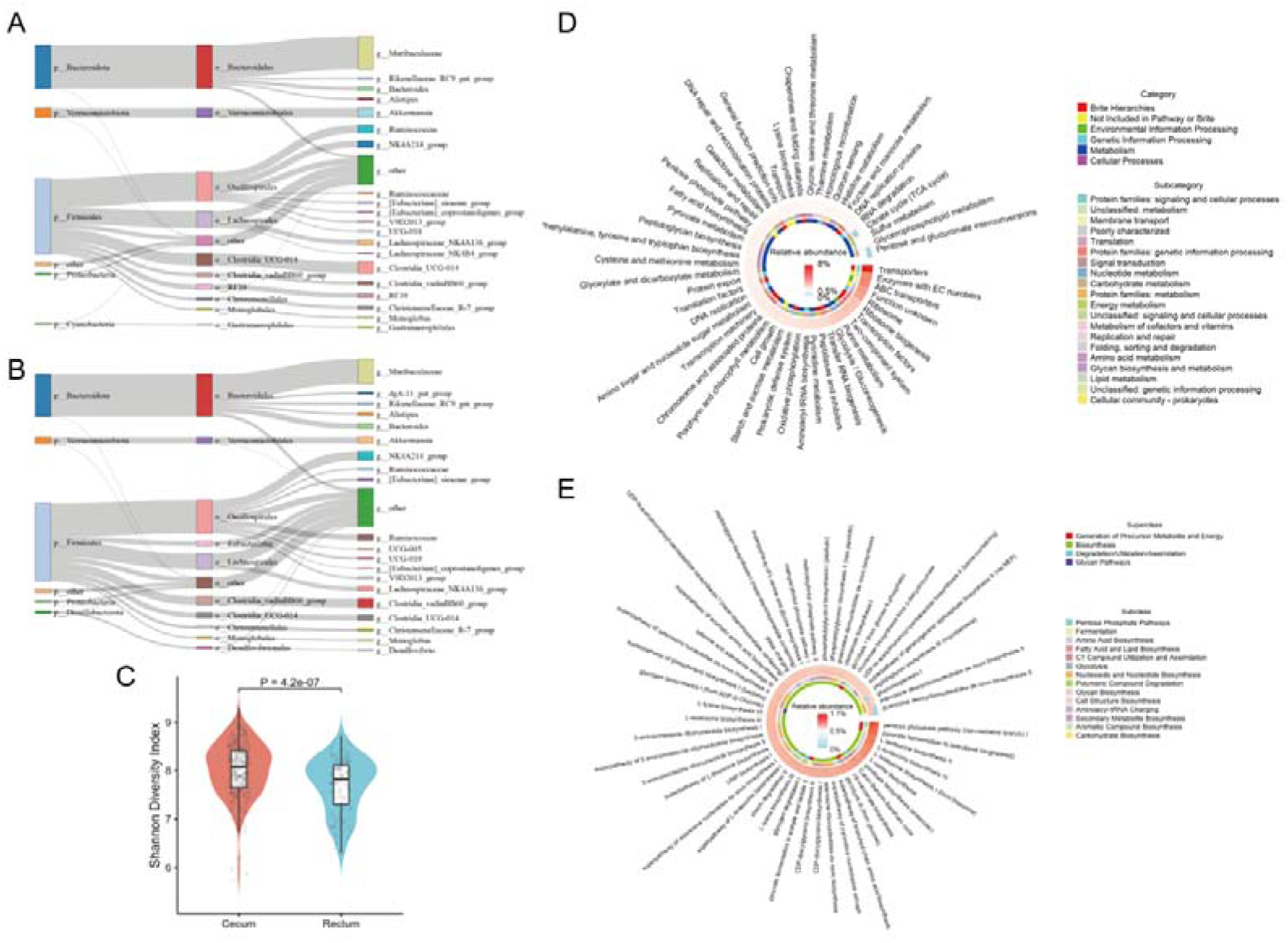
The microbial and functional landscape of rabbit cecal and rectal segments. A Composition and distribution of cecal microbiota from phylum to genus. The widths of the flows represent the relative abundance of each taxon. The top 5 phyla, top 10 orders, and top 20 genera are visualized. B Composition and distribution of rectum microbiota from phylum to genus. The widths of the flows represent the relative abundance of each taxon. The top 5 phyla, top 10 orders, and top 20 genera are visualized. C Comparison of Shannon diversity index between cecum and rectum samples. Statistical significance was determined using the unpaired Student’s t-test. D Prediction of microbial function based on the top 50 KEGG pathways.The outermost ring is the cecum, the second ring is the rectum. E Prediction of microbial function based on the top 50 MetaCyc pathways. The outermost ring is the cecum, and the second ring is the rectum.

The top 50 KEGG pathways in the cecum and rectum were almost identical (49/50), with the majority being related to metabolism (Fig. 1D, Table S7). In the top 50 MetaCyc pathways, the most enriched pathway in both the cecum and rectum was pentose phosphate pathway. This pathway is a common glycolytic pathway used by most organisms for the breakdown of glucose and related sugars, which can be considered as both a catabolic and anabolic pathway for energy production. (Fig. 1E, Table S8).

### 3.3. Host genetics and cecal microbiota are associated withvariation in loin muscle weight

Heritability (h²) represents the extent of host genetics influence on phenotypes, while the contribution of microbiota to the phenotypic variance is reflected by microbiability (m²). For LM, the heritability was estimated at 0.39. The microbiability of LM in the cecum was 0.13. These estimates indicated that host genetics predominantly governed LM, with the cecal microbiome exerting a moderate auxiliary influence. This substantial cecal m² estimate justifies further investigation into specific microbial taxa associated with this trait (Fig. 2A).

**Fig 2.**
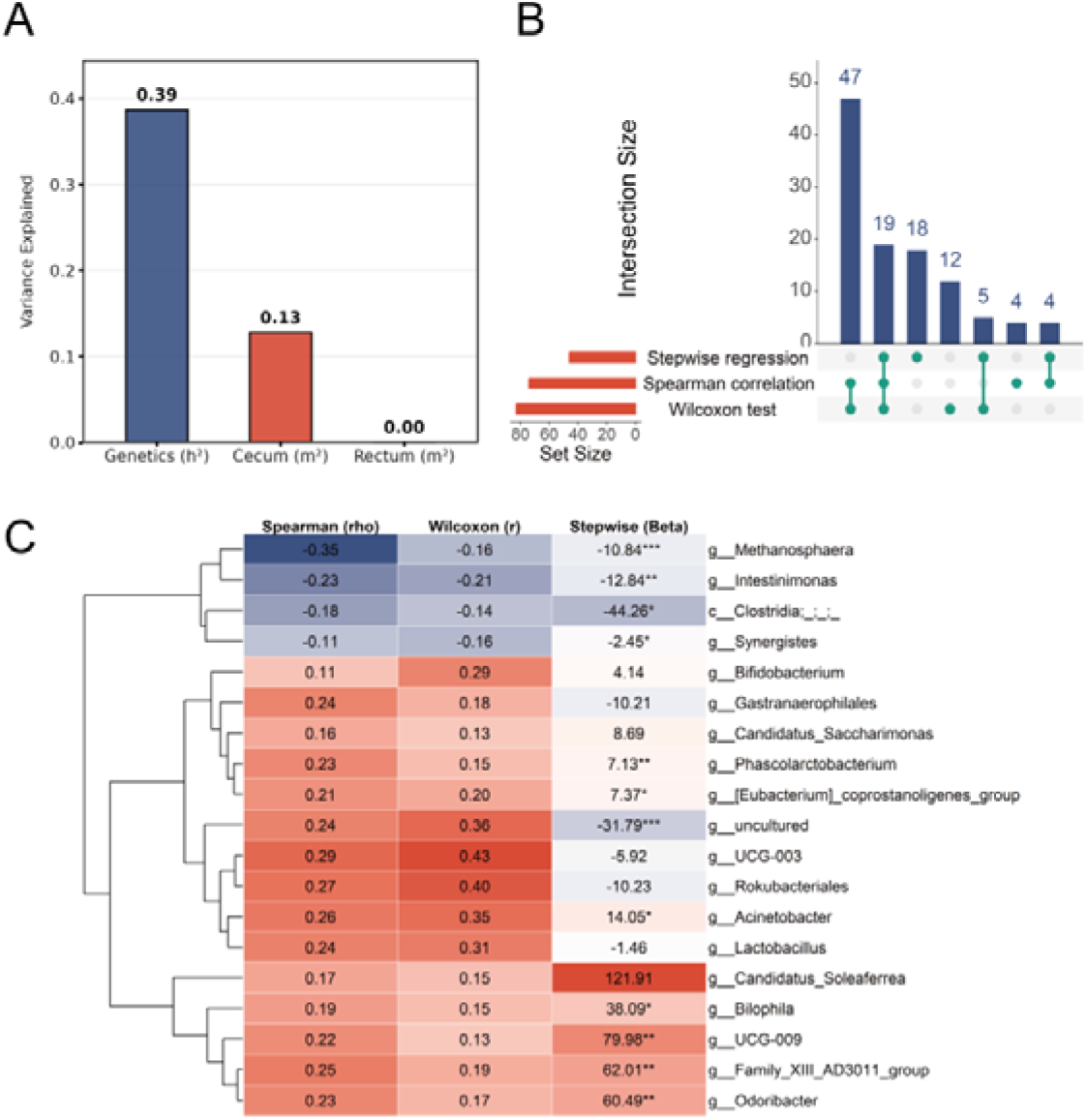
Genetic and microbial contributions to loin muscle and identification of key microbial drivers. A Heritability (*h^2^*) and microbiability (*m^2^*) of cecum and rectum for LM. B UpSet plots displaying the intersection of candidate genera associated with LM identified by three methods. C Heatmaps showing the association statistics for the high-confidence genera of LM, respectively. Stepwise regression significance levels: * *P* < 0.05, ** *P* < 0.01, *** *P* < 0.001.

To identify the specific bacterial taxa underlying these microbiability estimates, we employed a comprehensive screening strategy combining Spearman correlation, stepwise regression, and Wilcoxon rank-sum tests to narrow down the candidate taxa to those supported by multiple statistical approaches. Based on the intersection of these three strategies, we identified 19 high-confidence genera associated with LM (Fig. 2B, Table S9-11).

Among these candidates, the archaeal genus *Methanosphaera* exhibited the strongest negative correlation with LM (ρ = −0.35), while *Odoribacter* (ρ = 0.23) and the Family XIII AD3011 group (ρ = 0.25) emerged as strong positive drivers (Fig. 2C, Table S9).

### 3.4. Host genetic associations with microbial taxa differ between cecal and rectal segments

To explore whether host genetics were related to the structure of the gut microbiota, the correlation between the GRM and MRM of cecum and rectum were calculated using the Mantel test. Interestingly, a significant positive correlation was observed between the GRM and the Cecal MRM (r = 0.020, *P* = 3 × 10^-4^), indicating that a weak but statistically significant correlation was observed between host genetic relatedness and cecal microbial similarity, whereas no such global association was detected for the rectal microbiome(between the GRM and the rectal MRM r = −0.007, *P* = 0.69). Furthermore, the cecal MRM and rectal MRM showed no significant correlation (r = 0.008, *P* = 0.20), highlighting the distinct microbial architectures between these two gut segments.

In the cecum, mGWAS revealed that host genetic loci were significantly associated with 6 microbial genera and estimating their SNP-based *h^2^* (Fig 3, Table S12) The most significant association was observed for *Methanobrevibacter*, with the lead SNP located on Chromosome 5 (5:88203070, *P* = 3.11 × 10^-8^). In our prior phenotypic analysis, both Spearman correlation and Wilcoxon rank-sum tests confirmed that cecal *Methanobrevibacter* abundance was significantly associated with LM.

**Fig 3.**
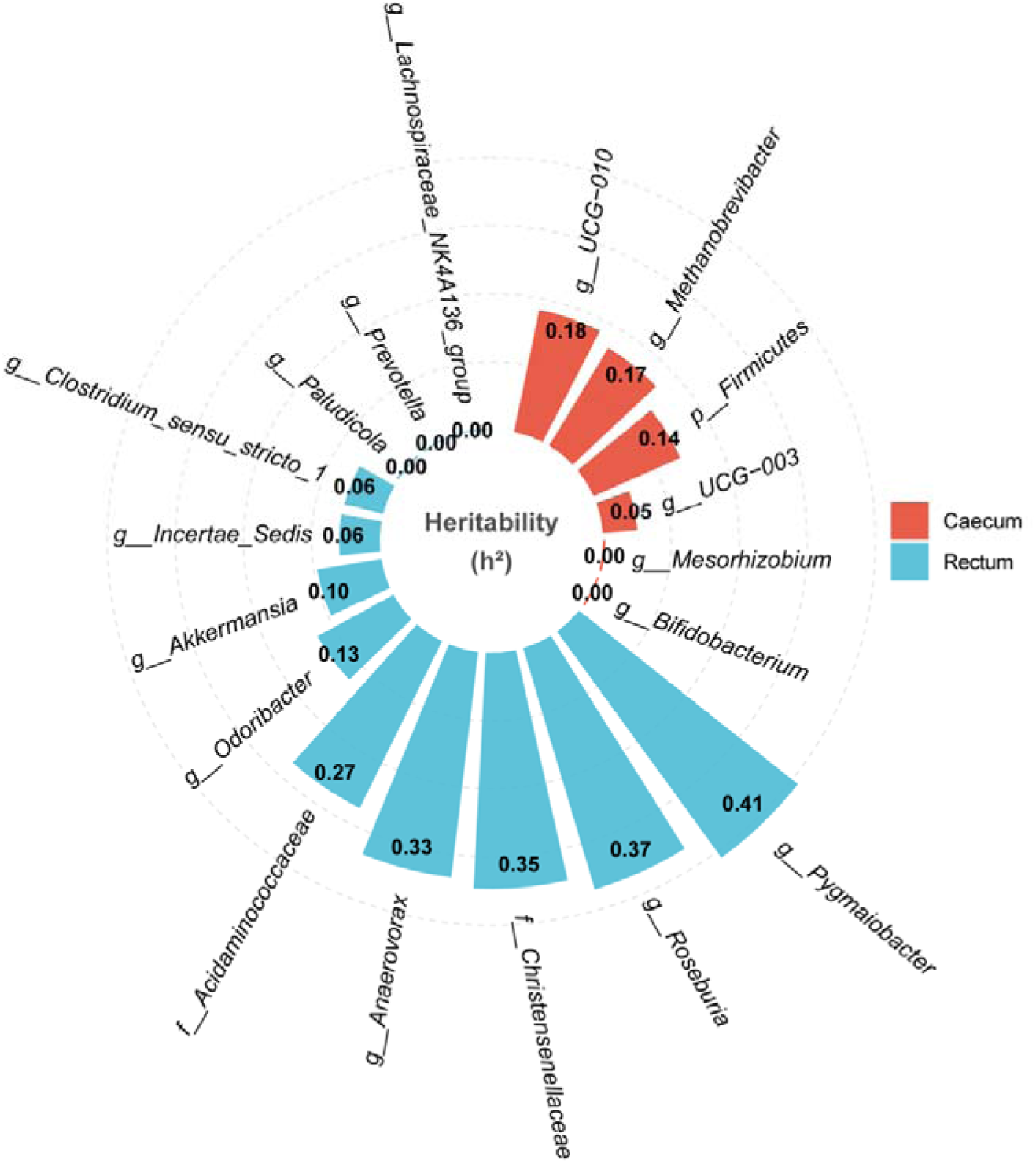
Heritability of gut microbiota in cecum and rectum

In the rectal content samples, while the global Mantel test failed to show statistical significance, mGWAS revealed more significant host-genetic associations for individual microbial genera, identifying 12 significant microbial genera and estimating their SNP-based *h^2^* (Fig 3, Table S12). These taxa included *Roseburia*, *Lachnospiraceae_NK4A136_group*, *Clostridium_sensu_stricto_1*, *Prevotella*, *Acidaminococcaceae*, uncultured genera from *Christensenellaceae*, *Pygmaiobacter*, *Akkermansia*, *Paludicola*, Odoribacter, *Anaerovorax*, and *Incertae_Sedis*. *Odoribacter* was a candidate genus associated with LM in our multi-strategy screening, it showed a highly significant association with a locus on Chromosome 20 (*P* = 5.61 × 10^-9^), supported by 302 SNPs. These findings outlined a putative “Host Genome – Microbiome – Phenotype” axis.

### 3.5. Cecal transcriptome provides gene-expression mediators linking host genetics *with loin muscle weight*

SRC were calculated to determine the relationship between cecum gene expression and phenotypic trait. The analysis identified 912 genes significantly correlated with LM, including 691 positively and 221 negatively correlated genes (P_adj < 0.05, Table S13). We identified DEGs between individuals ranking in the top and bottom 20% of the phenotypic traits groups. The results showed that 39 DEGs were associated with LM (|log_2_ FC| > 1, *P* < 0.05, Fig 4A, Table S14).

**Fig 4.**
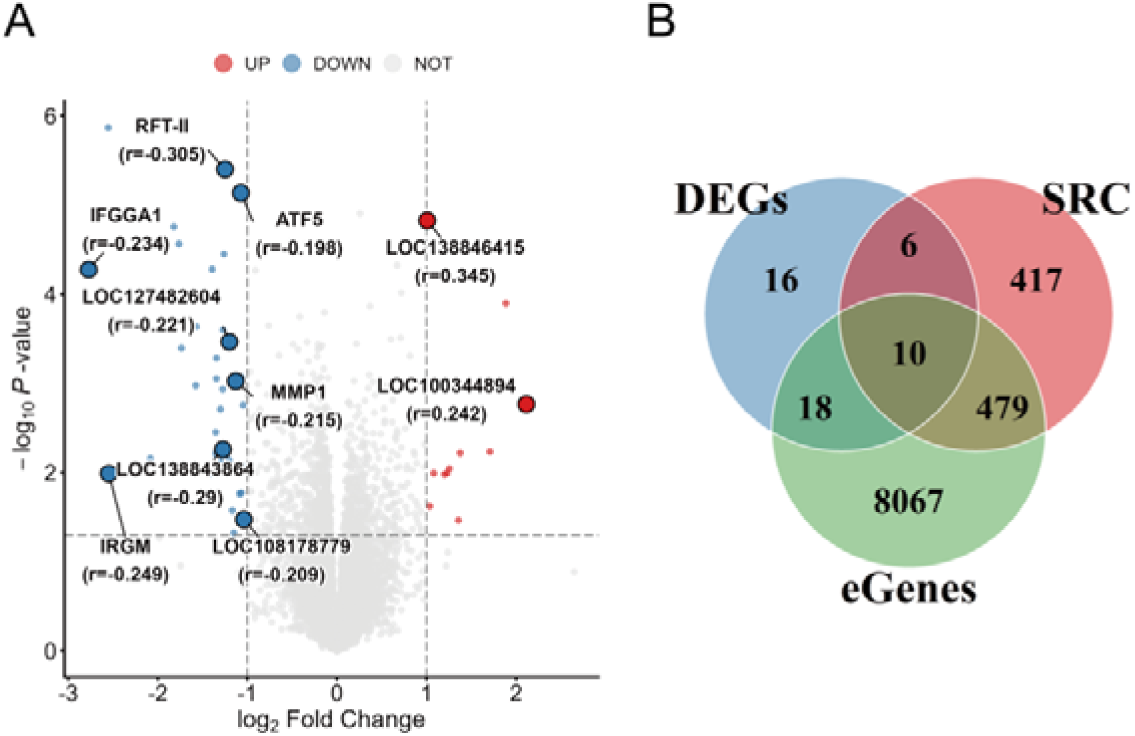
Identification of core candidate genes associated with loin muscle via integrated transcriptomic and genetic analyses. A Volcano plots displaying the differentially expressed genes (DEGs) LM. The labeled points highlight the core candidate genes identified by the multi-omics intersection strategy. The text annotations display the gene id and its correlation coefficient (r) by SRC. B Venn diagrams illustrating the overlap among DEGs, phenotype-correlated genes (SRC), and *cis*-eQTL regulated genes (eGenes) for LM.

To further determine which gene expression was under host genetic control, we performed *cis*-eQTL (expression quantitative trait loci) mapping. We focused on the protein coding and lncRNA genes. After adjustment, we identified 8575 eGenes (Table S15). We identified 10 overlapping genes across the three analytical strategies for LM. (Fig 4B). Importantly, these genes were distinct from those detected by SMR analysis, indicating that this transcriptome-wide integration strategy identified a novel set of genetically regulated candidate genes that complement the SMR findings.

### 3.6. SMR analyses prioritize cecal and rectal host gene–microbial taxon associations

GWAS analysis identified two genomic regions with suggestive significance for LM on Chromosome 7 and Chromosome 10. On Chromosome 7, the lead SNP 7:137689090 was mapped directly within the intron of the *DLGAP3* gene which supports *DLGAP3* as the functional candidate. On Chromosome 10, Genomic annotation revealed that these significant variants were primarily distributed across the intronic and downstream regions of *TCF4*. The peak SNP 10:23119913 was identified 27 kb downstream of the *TCF4* gene. Furthermore, another signal cluster was identified within the same locus, with variants 10:27105436 and 10:27105943 positioned 25 kb upstream of *MEX3C*. (Table S16)

To identify genes whose expression levels causally influence the phenotypic traits, we performed SMR analysis by integrating the *cis*-eQTL data with the GWAS summary statistics for LM. We identified 2 genes showing significant SMR associations (*P*_SMR_FDR < 0.05), both genes passed the HEIDI test (P_HEIDI > 0.05), suggesting that the associations were driven by a single causal variant rather than linkage disequilibrium (Table S17).

The most significant candidate was *GJB3* (*P*_SMR = 1.43 × 10^-4^, P_HEIDI = 0.54). As illustrated in the regional association plot (Fig 5A), the *cis*-eQTL signal regulating *GJB3* expression colocalized with the GWAS signal for LM at the SNP 7:137838280. Individuals carrying the C allele exhibited significantly lower expression levels of *GJB3* compared to the A allele (Fig 5B). Conversely, the same C allele was associated with significantly higher LM (Fig 5C). Collectively, the negative SMR effect size (*b*_SMR = −14.18) and the allelic trends suggest the allele C downregulates *GJB3* expression, which in turn leads to an increase in LM.

**Fig 5.**
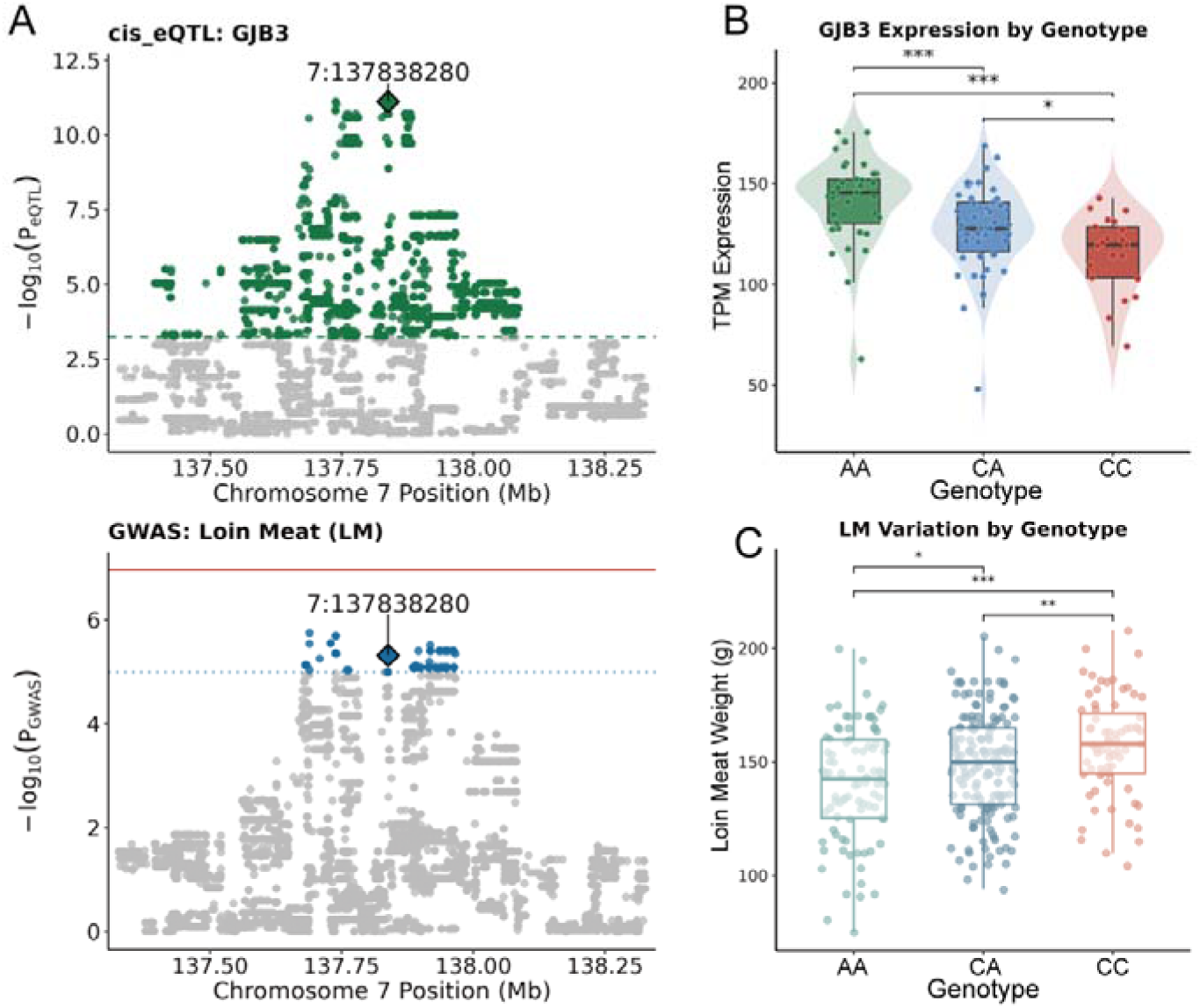
Causal association between *GJB3* expression and LM. A The top panel displays the *cis*-eQTL signals for *GJB3*. Green dots represent significant *cis*-eQTLs, grey dots represent non-significant. The bottom panel displays the GWAS signals for LM. The red line marks the significance threshold, and the dark blue line denotes the suggestive significance threshold. B, C Showing the association between SNP 7:137838280 genotypes with *GJB3* expression (B) and LM (C). Statistical significance was determined by the Wilcoxon test (* *P* < 0.05; ** *P* < 0.01; *** *P* < 0.001).

To identify genes whose expression levels influence the gut microbiota, we performed SMR analysis by integrating the *cis*-eQTL data with the mGWAS summary statistics from both cecal and rectal samples.

We identified a total of 14 significant host gene-microbiome pairs (cecum: 5, rectum: 9). 8 pairs (cecum: 3, rectum: 5) passed the HEIDI test. These results suggested that the associations were likely driven by a shared single causal variant. The remaining 6 pairs(cecum: 2, rectum: 4) showed significant SMR signals, but the HEIDI test was not applicable due to the limited number of instrumental SNPs. (Table S18) These associations likely represented effects driven by a single strong QTL.

In the cecum, SMR analysis prioritized a prominent host gene-microbial taxon association between *MRAP2* expression and the abundance of the archaeal genus *Methanobrevibacter* at the SNP 5:88261646(Fig. 6A). Individuals carrying the G allele exhibited significantly higher expression levels of *MRAP2* compared to the A allele (Fig. 6B). Conversely, the same G allele was associated with significantly lower relative abundance of *Methanobrevibacter* (Fig6.C). Consistent with the negative SMR effect size (*b*_SMR = −0.16), these allelic patterns suggest that genetically increased MRAP2 expression may be associated with reduced *Methanobrevibacter* abundance. This MRAP2-Methanobrevibacter association represents the main host-regulated microbial signal within the cecum-centered axis.

**Fig 6.**
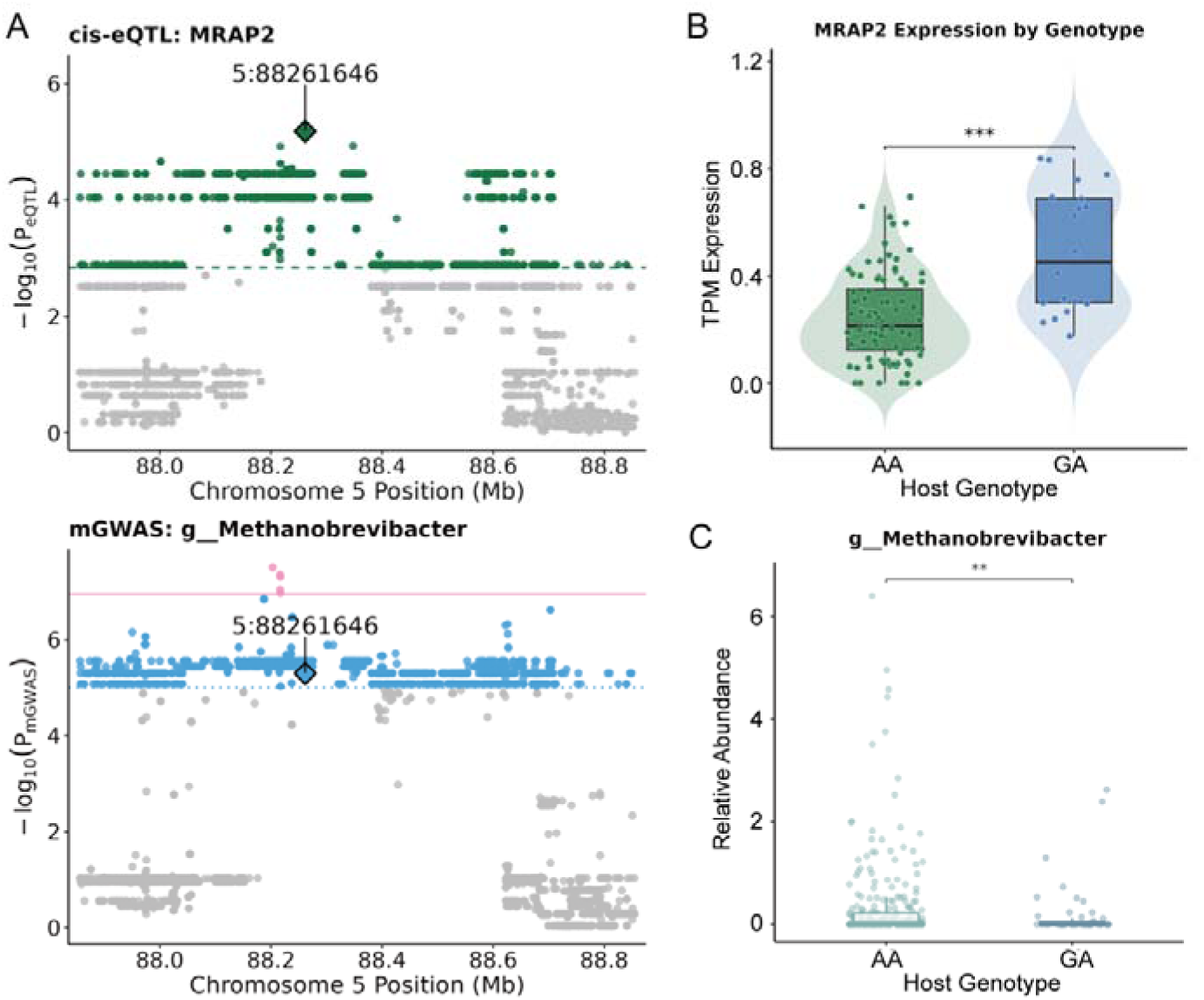
Causal association between host *MRAP2* expression and *Methanobrevibacter* abundance. A The top panel displays the *cis*-eQTL signals for *MRAP2*. Green dots represent significant *cis*-eQTLs, grey dots represent non-significant. The bottom panel displays the GWAS signals for *Methanobrevibacter*. The pink line marks the significance threshold, and the blue line denotes the suggestive significance threshold. B, C Showing the association between SNP 5:88261646 genotypes with *MRAP2* expression (B) and *Methanobrevibacter* (C). Statistical significance was determined by the Wilcoxon test (* *P* < 0.05; ** *P* < 0.01; *** *P* < 0.001).

For the rectal microbiome, SMR analysis identified a distinct segment-specific host gene–microbial taxon association between *SULF1* expression and the abundance of the butyrate-producing genus *Roseburia*, centered at SNP 6:94772750 (Fig. 7A). Individuals carrying the C allele exhibited significantly higher expression levels of *SULF1* than those with the T allele (Fig. 7B). Consistent with the positive SMR effect size (*b*_SMR = 0.82), these allelic trends suggest that genetically increased *SULF1* expression may be linked to increased Roseburia abundance in the rectal microbiome. This *SULF1*-Roseburia association supports the value of rectal microbiome data as a downstream comparative layer and suggests that host genetic associations with microbial taxa may show hindgut segment-specific patterns.

**Fig 7.**
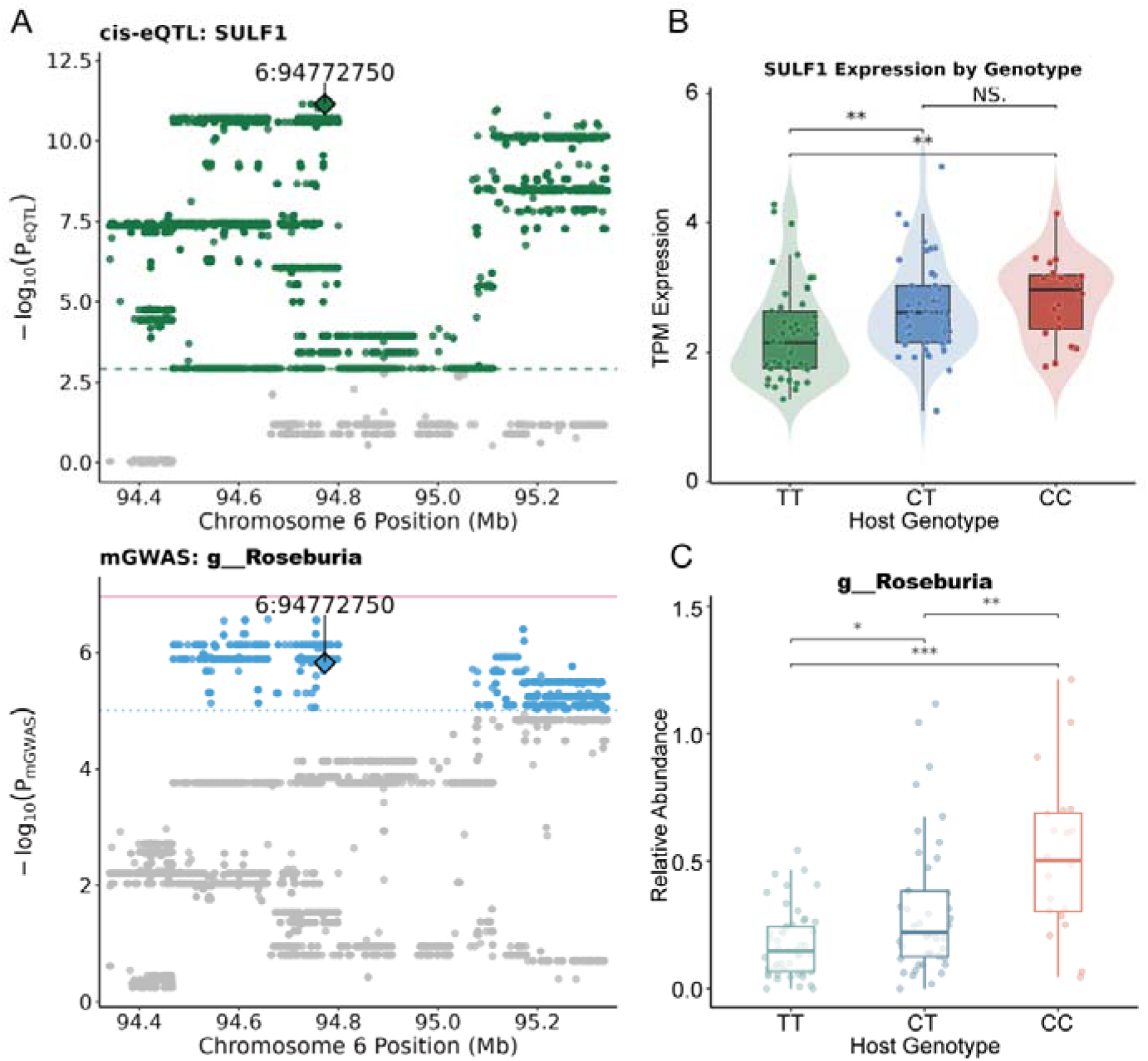
Causal association between host *SULF1* expression and *Roseburia* abundance. A The top panel displays the *cis*-eQTL signals for *SULF1*. Green dots represent significant *cis*-eQTLs, grey dots represent non-significant. The bottom panel displays the GWAS signals for *Roseburia*. The pink line marks the significance threshold, and the blue line denotes the suggestive significance threshold. B, C Showing the association between SNP 6:94772750 genotypes with *SULF1* expression (B) and *Roseburia* (C). Statistical significance was determined by the Wilcoxon test (* *P* < 0.05; ** *P* < 0.01; *** *P* < 0.001).

## 4. Discussion

The cecum serves as the primary fermentation site in rabbits, hosting a complex microbiota that is crucial for fiber digestion, fnutrient utilization, feed efficiency, and overall growth performance (Gidenne, 2015). Although previous studies have established that the rabbit gut microbiota is heritable (Velasco-Galilea et al., 2022), the mechanisms linking cecal microbiota to muscle deposition remain largely unclear. In our research, Firmicutes and Bacteroidota dominated the microbial communities. which aligns with previous study and its roles in carbohydrate degradation within hindgut fermenters (Combes et al., 2017; Stojanov et al., 2020). At the genus level, the robust colonization of glycan-degrading *Muribaculaceae* (Ormerod et al., 2016) and fiber-responsive Ruminococcaceae (Opdahl et al., 2018) further reflects the host’s evolutionary adaptation to a herbivorous diet and its reliance on microbial fermentation for energy harvest. We also noted the moderate presence of the sulfur-metabolizing Desulfobacterota (Hu et al., 2021; Liu et al., 2022a), potentially reflecting specific environmental or dietary exposures. Despite these localized taxonomic variations, functional predictions revealed a highly conserved metabolic core between the cecum and rectum. These findings support the use of the rectal microbiome as a downstream comparative segment, allowing us to distinguish cecum-centered host-microbial associations from broader segment-dependent patterns along the hindgut.

To comprehensively dissect the contributions of host genetics and gut microbiota to complex traits, our estimations of *h^2^* and *m^2^* revealed a distinct pattern of host dominance. Host genetics explained a substantial proportion of the phenotypic variance, whereas the cecal microbiome explained a smaller but detectable proportion of phenotypic variation. By contrast, the rectal microbiome showed a much weaker contribution, supporting the use of the rectum mainly as a comparative segment. These results align with findings in pigs (Camarinha-Silva et al., 2017), suggesting that growth is primarily driven by the host genome, with the gut microbiota serving as a significant auxiliary factor that tunes these phenotypes.

Consistent with the high heritability estimate for LM, our GWAS identified *MEX3C* on chromosome 10. This gene encodes a critical regulator of postnatal growth that post-transcriptionally enhances the translational efficiency of IGF1 (Jiao et al., 2012a). Furthermore, it acts as a pivotal metabolic regulator, directing energy resources to support skeletal muscle accretion (Jiao et al., 2012b).. On chromosome 7, we identified *DLGAP3*. While direct associations between *DLGAP3* and muscle traits are limited in the literature, our complementary SMR analysis in this region revealed a negative putative causal association between cecal *GJB3* expression and LM. Gap junctions are essential for intercellular communication. Based on the role of GJB3 in regulating cellular differentiation, lower GJB3 expression may facilitate intestinal epithelial maturation and nutrient absorption (Kibschull et al., 2014). This improved intestinal function could enhance nutrient availability for skeletal muscle growth.

The Mantel test identified a statistically significant positive correlation between host genetics and the cecal microbiota. This weak effect size aligns with the findings in human and chickens (Goodrich et al., 2016a; Goodrich et al., 2014; Rothschild et al., 2018; Xie et al., 2016). A weak or absent global correlation does not preclude genetic control over specific microbial groups. Our mGWAS analysis revealed that individual bacterial genera still exhibit significant and specific host genetic association signals. Consistent with previous reports in humans that the majority of the heritable taxa belongs to Firmicutes and Proteobacteria (Chen et al., 2018b; Davenport et al., 2015; Goodrich et al., 2016b). Furthermore, the abundance of Clostridia also identified significant host genetic variants of sheep, pigs, and chickens (Han et al., 2025; Li et al., 2024; Mani et al., 2022). In agreement with these findings, over half of the significant taxa in our mGWAS belonged to the class Clostridia (10/18). As major fermentative bacteria, Clostridia contribute to the degradation of complex carbohydrates and the production of metabolites that support host energy metabolism (Flint et al., 2012). The enrichment of host-selected Clostridia may therefore enhance nutrient utilization and energy harvest, ultimately contributing to muscle deposition.

The SMR analysis in this study found that host SNP 5:88261646_A_G reduces the abundance of the archaeon *Methanobrevibacter* by upregulating the expression level of *MRAP2* in cecal tissue. Furthermore, correlation analysis revealed that this methanogen was negatively correlated with LM. Although *MRAP2* is often described as a central appetite regulator in the hypothalamus (Asai et al., 2013), our finding of its expression in cecal tissue aligns with recent evidence regarding its peripheral function. Previous studies have also proven that *MRAP2* is expressed in human peripheral tissues (including the gut and pancreas), and MRAP2 is involved in the regulation of G-protein-coupled receptors including the growth hormone secretagogue receptor 1a (GHSR1a), namely the ghrelin receptor (Baron et al., 2019). *GHSR1a* has been confirmed to be expressed in the intestines of cattle, pigs, and chickens (Guan et al., 2025; Liu et al., 2022b; Teng et al., 2024). MRAP2 enhances GHSR1a signaling efficacy by blocking β-arrestin recruitment, thereby amplifying ghrelin signaling and subsequently stimulating gastrointestinal motility (Srisai et al., 2017). Intestinal transit time is a major determinant of microbiome structure. Compared with other commensal bacteria, *Methanobrevibacter* has a slower growth rate (Vandeputte et al., 2016); while it flourishes under conditions of slow transit to maximize caloric extraction (Armougom et al., 2009), the enhanced motility induced by *MRAP2* may leads to the mechanical wash-out of these slow-growing archaea, preventing their stable colonization. This explains the negative correlation we observed between *MRAP2* expression and *Methanobrevibacter* abundance. The association between *MRAP2*-mediated *Methanobrevibacter* reduction and LM increase represents a superior energy partitioning strategy under host nutrient-sufficient conditions. The inhibition of methanogenesis process shifts fermentation products from acetate to propionate (Ahvanooei et al., 2024; Castaneda et al., 2025). Propionate is a critical gluconeogenic substrate in the liver, providing glucose fuel essential for skeletal muscle growth, highlighting a superior host-driven energy partitioning strategy under nutrient-sufficient conditions. Collectively, these findings suggest that *MRAP2* may represent an important molecular link between host genetic variation and the cecal microbial ecosystem. Through its association with *Methanobrevibacter* abundance, *MRAP2* could contribute to host differences in nutrient utilization and muscle deposition.

Although the cecum was the primary focus of this study, the rectal microbiome provided an important downstream comparison for evaluating segment-specific host–microbial associations. The absence of a significant global correlation between the GRM and rectal MRM suggested that the rectal microbiome was less strongly aligned with overall host genetic relatedness than the cecal microbiome. Nevertheless, rectal mGWAS and SMR analyses identified genus-specific host genetic association signals, including the SULF1-Roseburia association. SULF1 encodes an extracellular endosulfatase involved in modifying heparan sulfate proteoglycans, which participate in epithelial development and intestinal barrier-related signaling (Liu et al., 2023) (Takemura and Nakato, 2017). In this study, genetically regulated cecal SULF1 expression was associated with Roseburia abundance in rectal contents. Because Roseburia is commonly regarded as a butyrate-associated genus, this association may indicate a segment-specific host–microbial link along the hindgut. However, because rectal transcriptome data were not available, we cannot determine whether SULF1 directly regulates the rectal microenvironment. Therefore, the SULF1–Roseburia result should be interpreted as comparative evidence for hindgut segment specificity rather than as a direct local regulatory mechanism.

Together, these findings support a cecum-centered model in which host genetic variation may be linked to loin muscle deposition through cecal gene-expression regulation and fermentation-related cecal microbial taxa. The rectal microbiome provided a useful downstream comparison, showing that host genetic associations with microbial taxa may not be uniformly shared across hindgut segments but instead may display segment-specific patterns. Future studies integrating metagenomics, intestinal metabolomics, direct measurements of short-chain fatty acids and methane production, and functional validation experiments will be needed to test the proposed host gene-microbiome links and clarify their roles in nutrient utilization and muscle deposition.

## 5. Conclusions

This study identified a cecum-centered host genome-cecal transcriptome-microbial taxon axis associated with loin muscle deposition in meat rabbits. The main findings prioritized fermentation-related cecal microbial taxa and host-regulated gene-microbe links, particularly the MRAP2-Methanobrevibacter association. Rectal microbiome analyses provided downstream comparative evidence for segment-specific host genetic associations with microbial taxa, exemplified by the SULF1-Roseburia association. These candidate host and microbial targets provide a basis for future functional validation and microbiome-informed nutritional strategies aimed at improving carcass traits in meat rabbits.

## Acknowledgements

This work was supported by the Shandong Special Economic Animal Industry Technology System [SDAIT-21-02], Qingdao Science and Technology Benefiting People Demonstration Project [25-1-5-xdny-26-nsh]. The authors sincerely thank the Livestock and Poultry Molecular Breeding Team at the College of Animal Science and Technology, Shandong Agricultural University, for their valuable support and assistance during this study.

## Declaration of generative AI and AI-assisted technologies in the writing process

During the preparation of this work the authors used ChatGPT in order to improve the English phrasing and enhance the readability of the text. After using this tool/service, the author(s) reviewed and edited the content as needed and take(s) full responsibility for the content of the published article.

## Data availability

Whole-genome resequencing data are available in the NCBI Sequence Read Archive (SRA) and the corresponding BioProject accession number is PRJNA1469912, and RNA-Seq data are available in the SRA under accession PRJNA1472903. 16S rRNA sequencing data can be accessed on SRA under the accession numbers PRJNA1469941.

## Supplementary Information

### Additional file 1

Table S1. Descriptive statistics for host phenotypes. Table S2. Summary statistics of whole-genome resequencing. Table S3. Summary statistics of 16S rRNA gene sequencing. Table S4. Summary statistics of bulk RNA-seq data. Table. S5. Phyla with relative abundance >1%. Table S6. Genus with relative abundance >1%. Table S7. Top 50 KEGG pathways with the highest average relative abundance. Table S8. Top 50 MetaCyc pathway with the highest average relative abundance. Table S9. SRC analysis performed between the abundance of microbial taxa in Cecum and LM. Table S10. LM comparison between the top and bottom 20% abundance groups of cecal genus-level relative abundance (Wilcoxon rank-sum test). Table S11. Summary of stepwise linear regression models identifying key cecal microbial genera associated with LM. Table S12. Detailed information on the SNPs associated with loin meat (LM). Table S13. Detailed information on the SNPs associated with relative abundance. Table S14. SRC analysis between the expression of each gene and LM. Table S15. Differential expression genes detected between the top and bottom 20% LM-ranked rabbits. Table S16. Summary of significant *cis*-eQTLs in cecum tissue. Table S17. The SMR test based on the GWAS results for LM. Table S18. The SMR test based on the mGWAS results.

### Additional file2

**Fig. S1.**
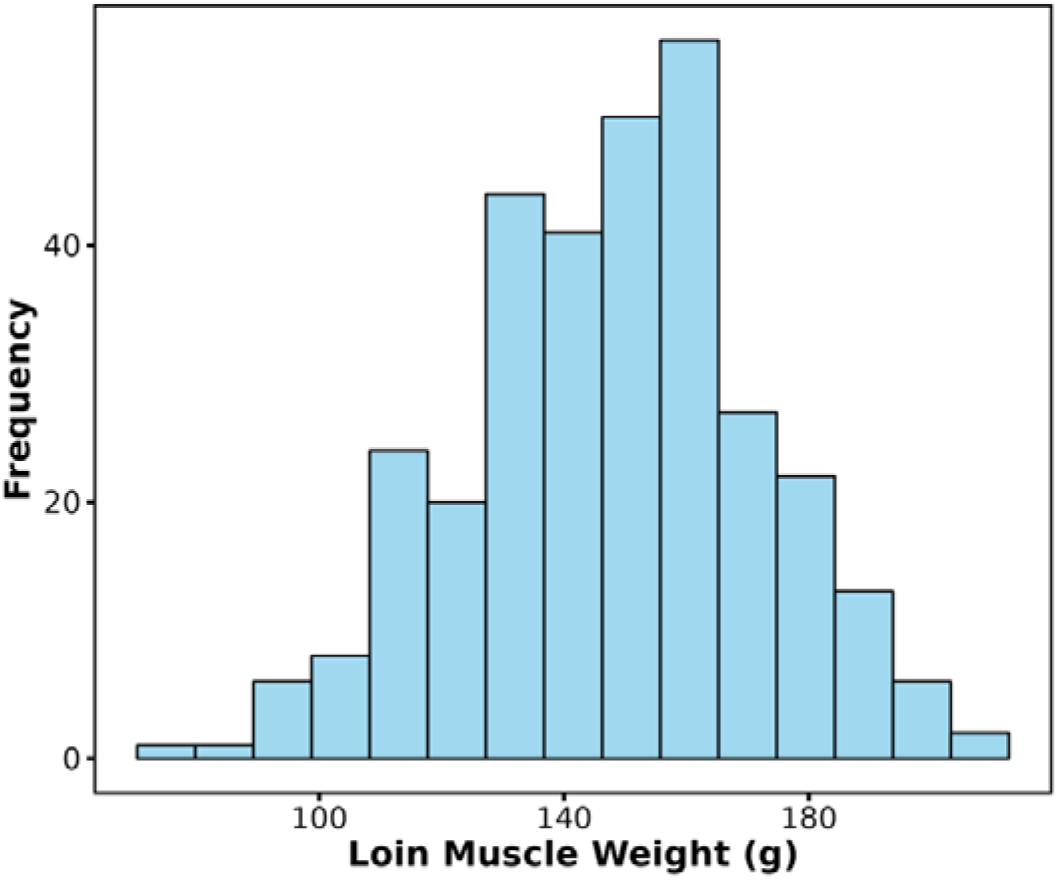
Phenotypic distribution of loin muscle (LM) weight.

### Additional file3

**Fig. S2.**
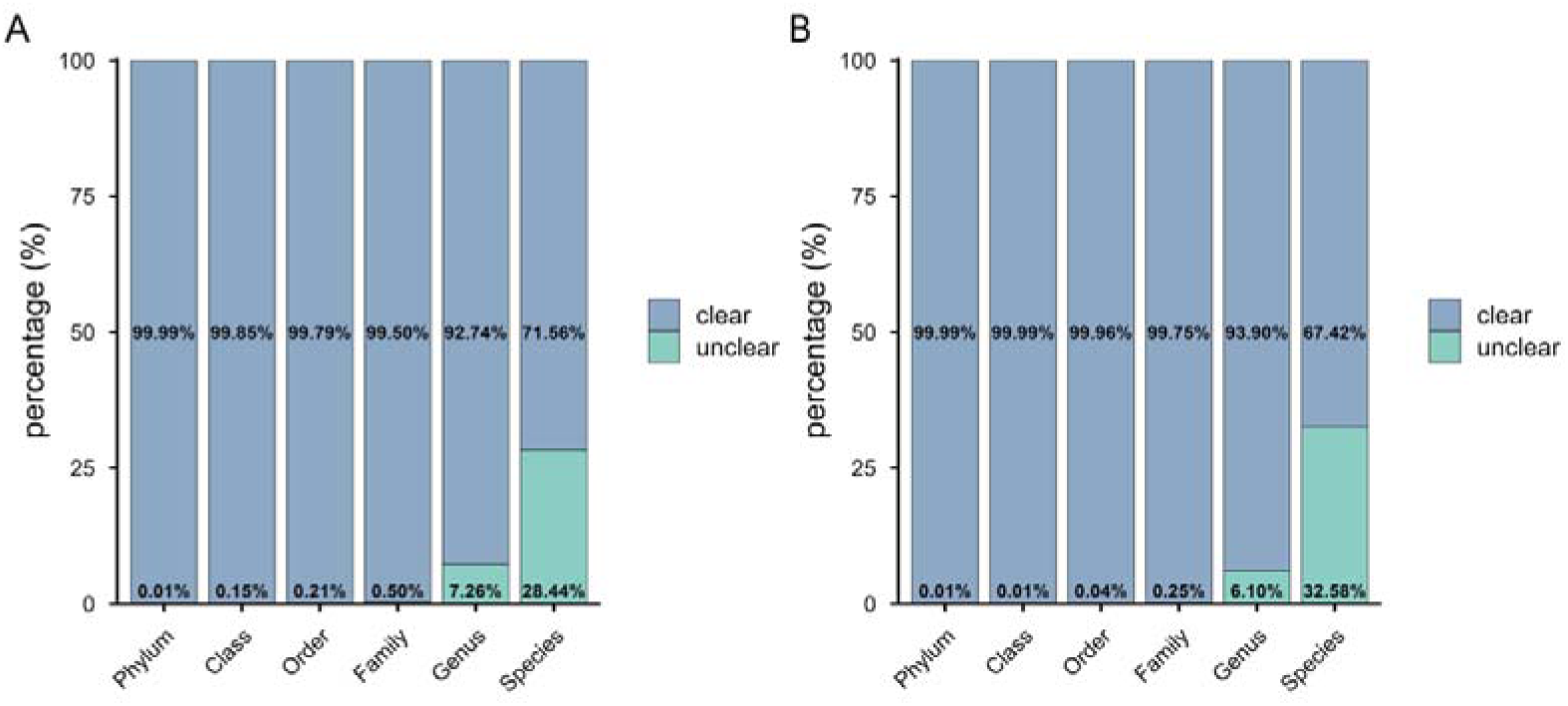
The proportion of reads can be accurately defined at all levels of classification. A: cecal, B: rectal.

